# Deciphering interaction fingerprints from protein molecular surfaces

**DOI:** 10.1101/606202

**Authors:** P Gainza, F Sverrisson, F Monti, E Rodolà, MM Bronstein, BE Correia

## Abstract

Predicting interactions between proteins and other biomolecules purely based on structure is an unsolved problem in biology. A high-level description of protein structure, the molecular surface, displays patterns of chemical and geometric features that *fingerprint* a protein’s modes of interactions with other biomolecules. We hypothesize that proteins performing similar interactions may share common fingerprints, independent of their evolutionary history. Fingerprints may be difficult to grasp by visual analysis but could be learned from large-scale datasets. We present *MaSIF*, a conceptual framework based on a new geometric deep learning method to capture fingerprints that are important for specific biomolecular interactions. We showcase MaSIF with three prediction challenges: protein pocket-ligand prediction, protein-protein interaction site prediction, and ultrafast scanning of protein surfaces for prediction of protein-protein complexes. We anticipate that our conceptual framework will lead to improvements in our understanding of protein function and design.

## Introduction

Interactions between proteins and other biomolecules are the basis of protein function in most biological processes. Predicting these interactions purely from protein structures remains one of the most important challenges in computational structural biology^1–4^. Many proteins bear evolutionarily conserved interaction signatures that can be effectively inferred from protein sequence and/or structure, and reveal their interactions with other biomolecules^5, 6^. Programs that exploit these signatures can be very effective at elucidating protein interactions and function^5, 7^, but these approaches rely on the existence of homologues from which to infer these interactions. The molecular surface^8^ is a higher-level description of protein structure that models a protein as a continuous shape with geometric and chemical features. We propose that molecular surfaces are *fingerprinted* with patterns of chemical and geometric features that reveal information about the protein’s interactions with other biomolecules. Our central hypothesis is that proteins with no sequence homology that undergo similar biomolecular interactions may display similar patterns, which may be difficult to grasp by visual analysis but could be learned from large-scale datasets. Here, we present MaSIF (Molecular Surface Interaction Fingerprinting), a general geometric deep learning^9^ method to recognize and decipher these patterns on protein surfaces, without explicit consideration of the underlying protein sequence or structural fold.

The molecular surface describes a protein structure based on solvent accessibility. It abstracts the underlying protein sequence, while displaying the geometric and chemical features that can enter in contact with other biomolecules. Its usefulness for many tasks involving protein interactions has long been known^7–9^, and has been the preferred structural description to study protein:solvent electrostatic interactions^10^. More recently, some efforts have captured molecular surface patterns with functional relevance, using techniques such as 3D Zernike descriptors^11–14^ and geometric invariant fingerprint descriptors (GIF)^15^. These approaches proposed ‘handcrafted’ descriptors, manually-optimized vectors which describe protein surface features. The limitation of these approaches is that it is hard, if not impossible, to tell a priori what are the right features, or combination of thereof, for a given prediction task.

In the field of computer vision, the emergence of deep learning techniques has led to an overwhelming trend of abandoning handcrafted features in favor of end-to-end data-driven task-specific features. Geometric deep learning^9^ (GDL) is a nascent field extending successful image-based deep neural network architectures, such as convolutional neural networks^16^, to geometric data such as surfaces, where these techniques have been shown to significantly outperform handcrafted feature extraction^17, 18^. MaSIF exploits GDL techniques to learn interaction fingerprints in protein molecular surfaces. First, MaSIF decomposes a surface into overlapping radial patches with a fixed geodesic radius (Fig. 1b), wherein each point is assigned an array of geometric and chemical features (Fig. 1c). MaSIF then computes a descriptor for each surface patch, a vector that encodes a description of the features present in the patch (Fig. 1d). Then, this descriptor can be processed in a set of additional layers depending on the application. The features encoded in each descriptor and the final output depend on the application-specific training data and the optimization objective, meaning that the same architecture can be repurposed for various tasks.

**Fig. 1.**
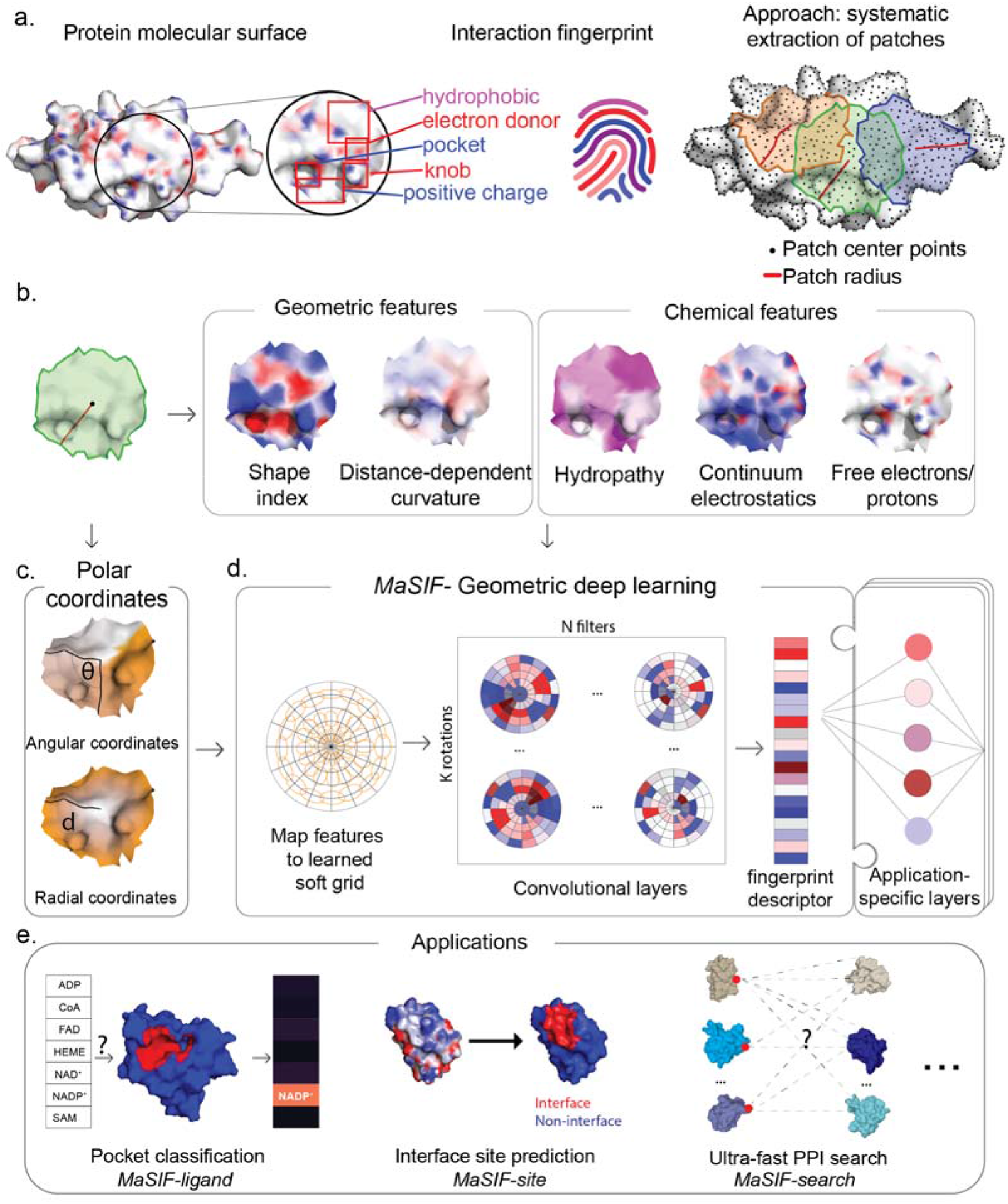
Overview of the MaSIF conceptual framework, implementation, and applications. **a**. Left, conceptual representation of a protein surface engraved with an interaction fingerprint, surface features that may reveal their potential biomolecular interactions. Right, surface segmentation into overlapping radial patches of a fixed geodesic radius used in MaSIF. **b**. The patches comprise geometric and chemical features mapped on the protein surface. **c**. Polar geodesic coordinates used to map the position of the features within the patch. **d**. MaSIF uses geometric deep learning tools to apply convolutional neural networks to the data. A fingerprint descriptor is computed for each patch and application-specific layers are then used to process the output. **e**. MaSIF is generalizable and applicable to multiple prediction tasks - a selected few are showcased in this paper.

To show the potential of the approach, we showcase three proof-of-concept applications (Fig. 1e): a) ligand prediction for protein binding pockets (*MaSIF-ligand*); b) protein-protein interaction (PPI) site prediction in protein surfaces, to predict which surface patches on a protein are more likely to interact with other proteins (*MaSIF-site*); c) ultrafast scanning of surfaces, where we use surface fingerprints from binding partners to predict the structural configuration of protein-protein complexes (*MaSIF-search*). Since our conceptual framework can be used in the absence of evolutionary information, it will be generally useful for biologists that search for interaction fingerprints in their proteins of interest. Importantly, the approach described in this paper represents a departure from learning on Euclidean structural representation and may enable the recognition of important structural features for protein function and design.

### MaSIF - A general framework to learn protein surface fingerprints

The conceptual and technical framework developed (MaSIF) is shown in Fig. 1 and described in detail in the Methods section. Briefly, from a protein structure we first compute a molecular surface discretized as a mesh according to the solvent excluded surface, and assign geometric and chemical features to every point (vertex) in the mesh (Fig. 1a-b). Then, we apply a geometric deep neural network to these features. The neural network consists of one or more *layers* applied sequentially; a key component of the architecture is the *geodesic convolution*, generalizing the classical convolution to surfaces and implemented as an operation on local patches^18^.

Around each vertex of the mesh, we extract a patch with geodesic radius of *r*=9 Å or *r*=12 Å (Fig. 1b, 1d). For each vertex within the patch, we compute two geometric features (shape index^19^ and distance-dependent curvature^15^) and three chemical features (hydropathy index^20^, continuum electrostatics^21^, and the location of free electrons and proton donors^22^), further described in Methods. The vertices within a patch are assigned geodesic polar coordinates (Fig. 1c): the radial coordinate, representing the geodesic distance to the center of the patch; and the angular coordinate, computed with respect to a random direction from the center of the patch, as the patch lacks a canonical orientation. In these coordinates, we then construct a family of learnable parametric kernels that locally average the vertex-wise patch features and produce an output of fixed dimension, which is correlated with a set of learnable filters^17^. We refer to this family of learnable parametric kernels as a *learned soft polar* grid. Note that since the angular coordinates were computed with respect to a random direction, it becomes essential to compute information that is invariant to different directions (*rotation invariance*, Fig. 1c). To this end, we perform *K* rotations on the patch and compute the maximum over all rotations^18^, producing the geodesic convolution output for the patch location. The procedure is repeated for different patch locations similarly to a sliding window operation on images, producing the surface *fingerprint* at each point, in the form of a vector that stores information about the surface patterns of the center point and its neighborhood. The learning procedure consists of finding the optimal parameter set of the local kernels and filter weights. The parameter set minimizes a cost function on the training dataset, which is specific to each application that we present here. We have thus created descriptors for surface patches that can be further processed in neural network architectures. Next, we will present various ways to leverage them to identify interaction fingerprints on protein surfaces.

### Molecular surface fingerprinting to classify ligand binding pockets

Interactions between proteins and metabolites play a fundamental role in cellular homeostasis, yet our knowledge of these interactions is extremely limited^23^. We propose that the interaction fingerprints in protein surfaces hold sufficient information to decipher the metabolite-binding preference of protein pockets. To test this, we developed *MaSIF-ligand*, a classifier that predicts the metabolite binding preference of a pocket. For this proof-of-concept we used seven important cofactors: ADP, NAD, NADP, FAD, SAM, CoA and HEME, metabolites with abundant structural data available.

We trained MaSIF-ligand on a large set of cofactor-binding proteins, where sequences were clustered to remove redundancy from the training and test sets (see Methods). The balanced accuracy on an independent test set was used to gauge the classification power of MaSIF-ligand. In this case with 7 cofactors, the expected balanced accuracy for a random classifier is 0.14. We first trained MaSIF-ligand with all features (geometry and chemistry) and obtained an accuracy of 0.78 and a balanced accuracy of 0.73. To investigate the importance of the features we limited the set of features to geometric or chemical features which reduced the balanced accuracy to 0.55 and 0.65, respectively (Fig. 2c).

**Fig. 2.**
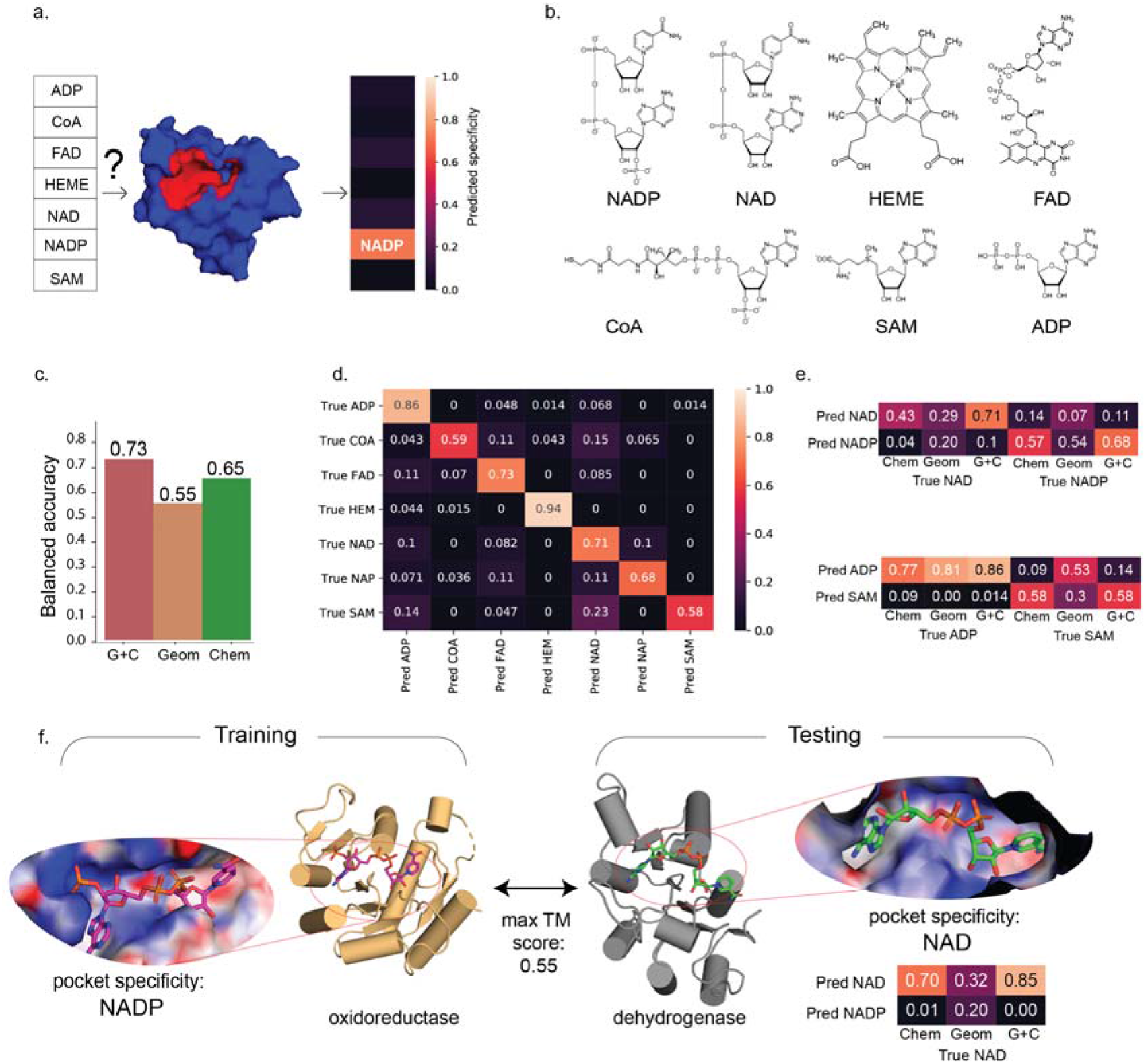
Classification of ligand binding sites using MaSiF-ligand. a. Schematic representation of the prediction task. The neural network receives a protein pocket as input and classifies it into seven categories to reflect the predicted binding preference. b. Structures of the seven cofactors that bind proteins considered for the prediction task. c. Balanced accuracy of the prediction of the specificity of binding sites using all features (G+C - Geometry and Chemistry), only geometric features (Geom), or only chemical features (Chem) d. Confusion matrix of ligand specificity on a MaSIF-lig trained with all features. e. Subset of the confusion matrices showing the importance of the features in distinguishing pockets between highly similar ligands. f. Specific example on a protein fold that recognizes two similar ligands and yet is correctly predicted. A bacterial dehydrogenase in the test set binds to NAD (PDB id: 2O4C)^24^, while its closest structural homologue in the training set corresponds to a mammalian oxidoreductase (PDB id: 2YJZ), which binds to NADP^25^.

The confusion matrix of MaSIF-ligand with all features (Fig. 2d) shows a variable performance across ligands, which is perhaps not surprising in the case of HEME (accuracy of 94%), as HEME is chemically dissimilar and larger compared to the other cofactors. More challenging is the distinction between similar ligands, namely in the analysis of the confusion data between two highly similar cofactors: SAM vs. ADP and NADP vs. NAD. In both cases, the geometric features are not sufficient for correct identification and it is mainly the chemical features which allow the network to correctly assign the pockets (Fig. 2e). The capacity of MaSIF-ligand to distinguish the features from cofactors with such a high structural and chemical similarity is remarkable, especially for NADP vs. NAD, where the only difference is a single phosphate group on the adenosine moiety. To understand these successful predictions, we analyzed the pocket features of an NAD-binding bacterial dehydrogenase^24^ in our test set and its closest structural homologue in the training set, a mammalian oxidoreductase which binds to NADP (Fig. 2f)^25^. For each of the two proteins, we analyzed the regions of the pocket that were giving the neural network the highest discrimination score between NAD vs. NADP. For this, we colored each point in the pocket based on its importance in discriminating between NAD and NADP (see Methods) (Supp. Fig. 1). The highlighted regions clearly show that the patches centered around the additional NADP phosphate in the oxidoreductase:NADP pocket determine its specificity for NADP vs. NAD, while in the dehydrogenase:NAD pocket, the region around the adenine moiety, near the location where NAD and NADP differ, are crucial to classify the pocket as NAD-binding. The prediction probabilities for the dehydrogenase:NAD pocket show a clear dependency on the chemical features (Fig. 2f, right). This is further confirmed by a visual inspection of the Poisson-Boltzmann electrostatics of the pocket, as it shows that the oxidoreductase:NADP pocket (Fig. 2f, left) has a stronger positive charge distribution according to the Poisson-Boltzmann electrostatics. This is consistent with its binding to the more negatively charged NADP. Despite the lack of global sequence homology between the test and training sets, *MaSIF* can decipher the surface interaction fingerprints to determine the binding preference of each pocket. As shown by the dehydrogenase:NAD example, even though proteins from the same structural family can bind to the same cofactor, MaSIF can infer the correct cofactor based purely on surface features, without explicit consideration of the underlying amino acids. A retrospective analysis shows that during the classification process, MaSIF accurately pinpoints critical sites that determine the specificity of NAD vs. NADP. Overall, the interaction fingerprints in protein surfaces could be an additional source of information available to biologists to infer important protein:ligand interactions.

### Predicting protein binding sites based on interaction fingerprints

In the previous section we focused on determining the specificity of a protein surface given highly similar ligands. A much broader classification is to infer an entire, broad, class of molecules based on the surface fingerprints. Inspired by previous work on PPI site prediction^26–28^, we developed MaSIF-site, a classifier that receives a protein surface as input and outputs a predicted score for each surface vertex on the likelihood of being involved in a PPI (Fig. 3 a.).

**Fig. 3.**
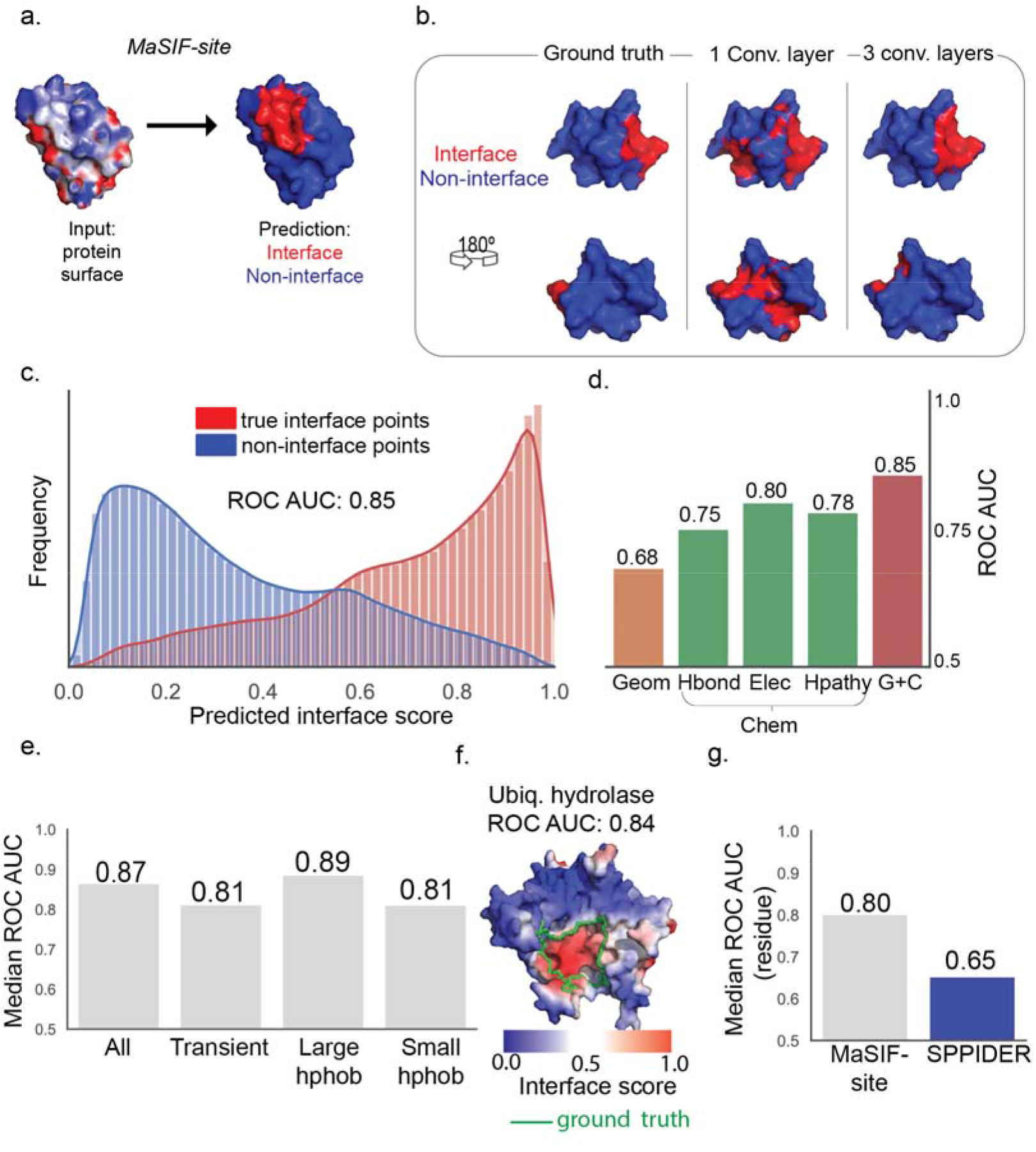
Prediction of surface patches involved in PPIs. **a**. Schematic representation of the interface site prediction workflow. The MaSIF-site receives as input a protein surface with a descriptor vector and outputs a surface score that reflects the predicted interface propensity (red for high interface propensity, blue for low propensity). **b**. Visual comparison between MaSIF-site with a network with 1 convolutional layer vs. 3 convolutional layers. **c**. Distribution of predicted scores for true positives (red) vs. true negatives (blue) for a network trained with all features. d. ROC AUC scores for ablation studies with networks trained with different subsets of features: only geometric (Geom), only the location of free electrons/proton donors (hbond), Poisson-Boltzmann electrostatics (elec), the hydropathy index (hpathy), and all features (G+C). **e. Left:** Median ROC AUCs (per protein) for selected subsets of proteins. **All** - full test set containing all proteins; Transient - proteins forming known transient interactions; **Large hphob** - protein complexes with interfaces composed of mostly hydrophobic residues; **Small hphob** - protein complexes with small hydrophobic interfaces. f. An illustrative example of a protein with an ROC AUC close to the median of 0.84, which is close to the median of MaSIF-site. e. Comparison of MaSIF-site with the SPPIDER predictor for the set of transient interactions. Results are shown as the median ROC AUC per protein, evaluated on a per-residue basis.

We tested MaSIF-site on a large dataset of protein structures, separated by both sequence and structural splits (see Methods). To evaluate the performance of our predictor, we use the receiver operator characteristic (ROC) area under the curve (AUC). We observed that this task greatly exploits the potential of deep learning approaches, since we can exploit multiple layers to build superior predictions (Fig. 3b). With just one convolutional layer the ROC AUC score for MaSIF-site reaches 0.77 (Supp. Fig. 2), shown in a specific example in Fig. 3b. Using three layers of geodesic convolution, MaSIF-site reaches a ROC AUC of 0.85. A distribution plot of all scores shows a strong separation between the predicted true and false interfaces (Fig. 3c). We investigated which surface features were the most important for the predictor’s performance by testing each feature set independently: using only geometric features (Geom), the ROC AUC score drops to 0.68; using only the location of free electrons and proton donors (hbond), ROC AUC=0.75; using only Poisson Boltzmann continuum electrostatics (elec), ROC AUC=0.80; and using only the hydropathy index of the vertices (hpathy), ROC AUC=0.78 (Fig. 3c).

Since surfaces involved in PPIs can be classified according to different biophysical (e.g. obligate vs transient) and structural/chemical (e.g. large vs small, hydrophobic vs polar, etc.) properties, we next asked whether MaSIF-site had a biased performance for a particular type of surface (Fig. 3d). For all interactions, the prediction accuracy reached a ROC AUC of 0.87, while for a subset of known transient interactions the ROC AUC was 0.82. We observed that proteins with large hydrophobic interfaces had a slightly better performance (ROC AUC: 0.89) than those with the smallest hydrophobic surfaces (ROC AUC: 0.82). The median AUC value is illustrated with the example of Ubiquitin Hydrolase protein, where the prediction reaches an AUC of 0.84, close to the median (Fig. 3f).

The prediction of PPI sites has been explored previously^26–28^ with predictors such as SPPIDER^27^ among the top performing and most used site predictors. We selected the subset of transient interactions, which are likely the more challenging to predict, and compared the performance of MaSIF-site with that of SPPIDER (see Methods). The median ROC AUC per protein, at the residue basis, is 0.80 for MaSIF-site, while it is 0.65 SPPIDER (Fig. 3g). Thus, we see a clear improvement in classification power with respect to an established predictor, which we further illustrate with a comparison between the two methods in four randomly chosen proteins from the transient test set (Supp. Fig. 3b).

Computational design efforts are making remarkable progresses in engineering de novo PPIs. The interaction sites in *de novo* proteins cannot be inferred from evolutionary information. We used MaSIF-site to predict three designed interfaces that have been experimentally validated: an influenza inhibitor^29^ (Fig. 4a), a homo-oligomeric cage protein^30^ (Fig. 4b), and an epitope-scaffold used as an immunogen^31^ (Fig. 4c). Each of these designed proteins was based on a wild type native protein with no binding activity, and in each case, we compared their surface interface score with that of the non-interacting wild type. We can observe a clear labeling of the interface only for the designs. Overall, exploring surface fingerprints with deep learning may be a powerful approach to identify the sites of interactions to other proteins that can guide experimental efforts for PPI validation, for paratope/epitope prediction, or small molecule binding sites, for cases where evolutionary or experimental information may not be available.

**Figure 4.**
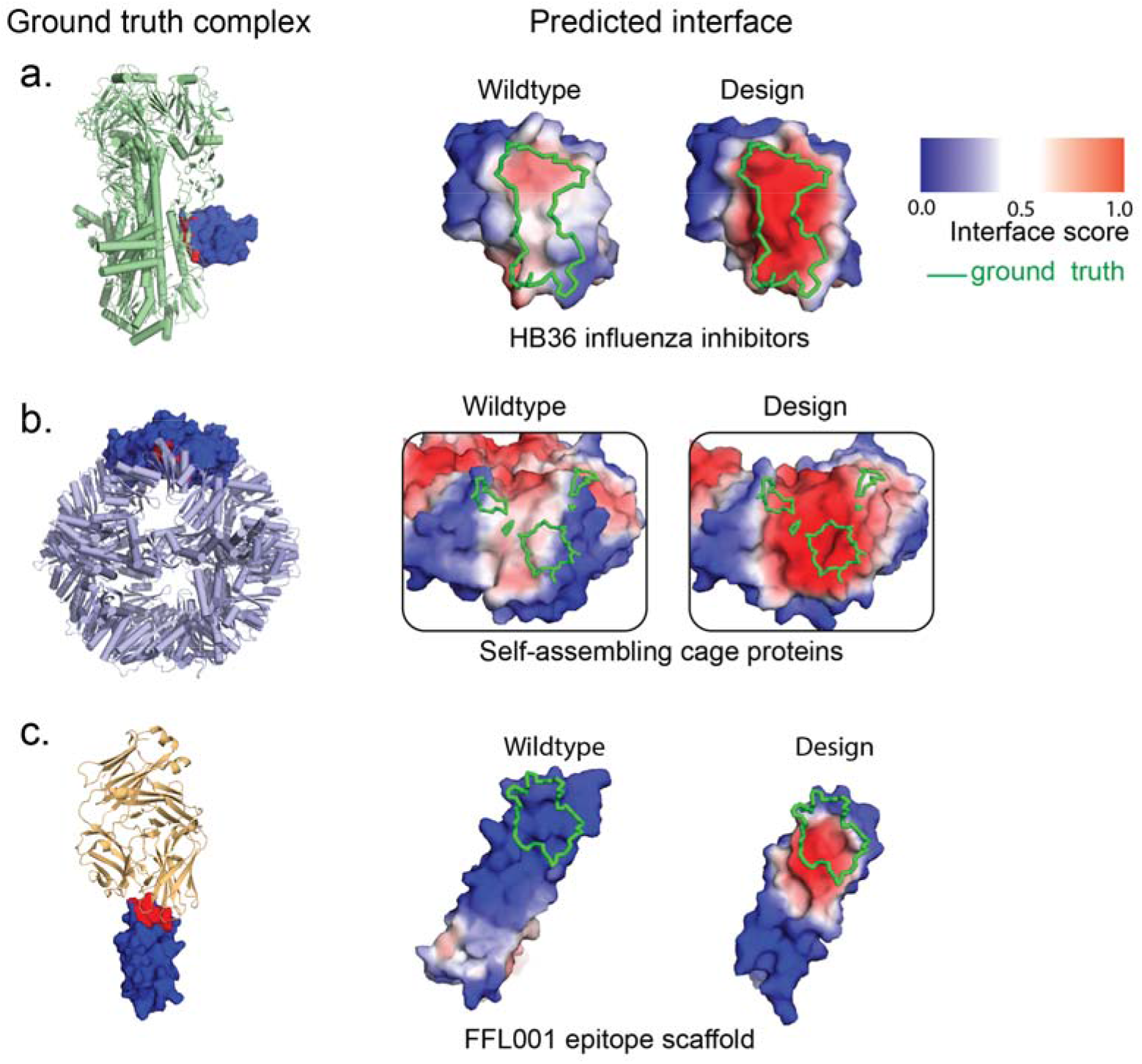
Prediction of PPI sites on a set of computationally designed proteins. **a**. Designed HB36 influenza inhibitors (PDB id: 3R2X) vs. the wild type scaffold protein (PDB id: 1U84). **b**. Designed self-assembling nanocage protein (PDB id: 3VCD) vs. the wild type scaffold (PDB id: 3N79) c. Designed Respiratory syncytial virus epitope-scaffold (PDB id: 4JLR) vs. wild type scaffold (PDB id: 1ISE). MaSIF-site was tested on a set of de novo computationally-designed proteins involved in PPIs, where the prediction on the designed binders can be compared to the corresponding native proteins. **Left**: crystal structures of the designed complexes, with the binder shown in surface representation, and the PPI buried surface area shown in red. **Middle**: prediction on the non-designed native protein scaffolds. **Right**: prediction on the designed proteins.

### Ultrafast scanning of interaction fingerprints for prediction of protein-protein complexes

In the previous two sections we exploited MaSIF to classify interaction fingerprints by the type of molecule they preferentially bind to. As a last example of MaSIF, we show how we can exploit MaSIF’s encoding of fingerprints as vectorized descriptors to predict specific interactions between proteins. This encoding, inspired by earlier work on GIF descriptors^15^, is attractive because, once the vectors are precomputed, nearest-neighbor techniques can scan billions of vectors in milliseconds^32^. The gain in computational cost at runtime enables broad structural searches across large databases, moving away from the paradigm of 1 binder vs. 1 target typical of docking programs, to one of many binders vs. many targets. This is important for tasks such as protein design, where docking tools are used to search for structural templates to use as starting points for the design of novel PPIs or ligand-binding proteins^29, 33^. Thus, we introduce *MaSIF-search* a new paradigm for the fast search of protein binding partners based on surface fingerprints, which is then complemented with a surface alignment stage to generate docked complexes.

MaSIF-search learns to identify the patterns that make two surface patches interact. We assume that proteins interact through surface patches with complementary geometric and chemical features (*complementary* fingerprints). To formalize this assumption, numerical features of one protein partner are inverted (multiplied by −1). The ultimate goal is that MaSIF-search will produce similar descriptors for pairs of interacting patches, and dissimilar descriptors for non-interacting patches (Fig. 5a). Thus, identifying potential binding partners is reduced to a comparison of numerical vectors.

**Figure 5.**
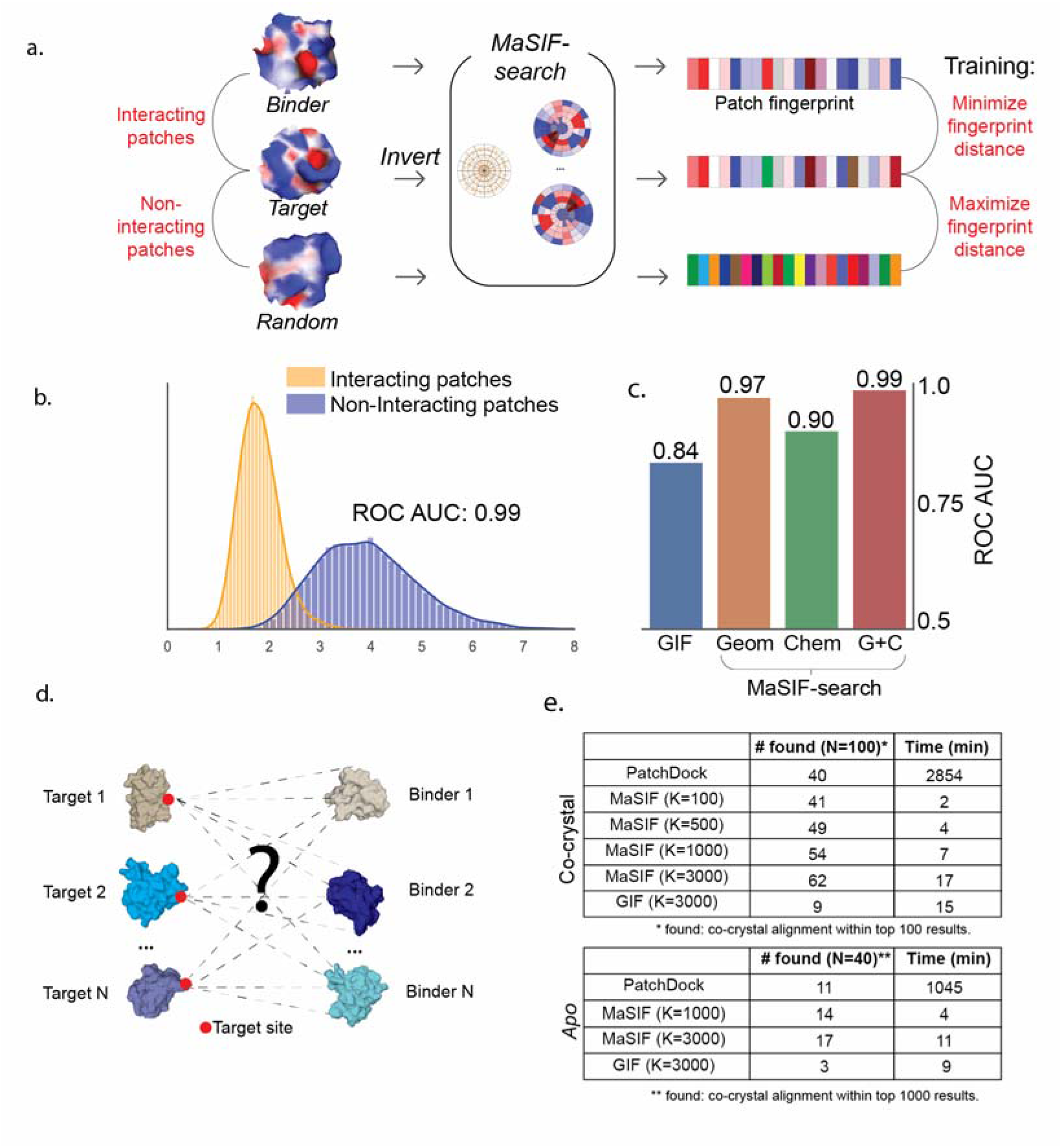
Prediction of PPIs based on surface fingerprints. **a**. Overview of the MaSIF-search neural network optimization to output fingerprint descriptors, such that the descriptors of interacting patches are similar, while those of non-interacting patches are dissimilar. The features of one of the interacting patches are symmetrized to enable the minimization of the fingerprint distance with the exception of the hydropathy features. **b**. Distribution of fingerprint distances showing interacting (yellow) and non-interacting (blue) patches for the test set. MaSIF-search was trained and tested on both geometric and chemical features. **c**. Comparison of the performance between different fingerprint features shown in ROC AUC. GIF - ROC AUC for GIF fingerprint descriptors^15^; Geom: MaSIF-search trained with only geometric features; Chem: MaSIF-search only with chemical features; S+C - geometry and chemistry features. **d**. We benchmarked MaSIF-search by performing a large-scale docking of N binder proteins to N known targets with site information. **e**. Results on the large scale docking benchmark for PatchDock, MaSIF-search (with multiple values of K) and GIF descriptors using our second stage method. **# solved:** number of target:binder complexes within 5 Å iRMSD found in either the top 100 (for bound cases) or top 1000 (for unbound cases). **Time:** Processing time in minutes for each algorithm - excludes pre-computation time for fingerprint-based algorithms.

To test this concept, we assembled a database with >100K pairs of interacting protein surface patches with high shape complementarity, as well as a set of randomly chosen surface patches, to be used as non-interacting patches. A trio of protein surface patches with the labels, *binder*, *target*, and *random* patches were fed into the network of MaSIF-search (Fig. 5a). The neural network is simultaneously trained to minimize the Euclidean distance between the fingerprint descriptors of *binders* and *targets*, while maximizing the dissimilarity between targets and random (see Methods).

Performances on the test set show that the descriptor Euclidean distances for interacting surface patches is much lower than that of non-interacting patches, resulting in a ROC AUC of 0.99 (Fig. 5b). Our method is directly comparable to the previously proposed handcrafted GIF descriptors^15^, which were proposed for a similar application: screening functional surfaces. Tested on our test set, GIF descriptors show a ROC AUC of 0.84, significantly lower than that of MaSIF-search (Fig. 5c). Testing MaSIF-search using only chemical or geometric features, we obtained ROC AUCs of 0.90 and 0.97, respectively. It is remarkable that chemical features alone can provide such a high discriminative power, and the improvement from 0.97 to 0.99 is a highly significant one. We observed that MaSIF-search has superior performance on high shape complementarity PPIs, as training/testing on interacting patches with lower shape complementarity results in lower performance (Supp. Fig. 6).

Next, we used MaSIF to predict the structure of known protein-protein complexes using a surface fingerprint-based search followed by structural alignment of surface patches. Briefly, the MaSIF-search workflow entails two stages: I) scanning a large database of descriptors of potential binders and selecting the top K by descriptor similarity; II) three-dimensional alignment of the complexes exploiting fingerprint descriptors of surrounding points, followed by a reranking of predictions according to fingerprint descriptors (see Methods and Supp. Fig. 4).

We benchmarked MaSIF-search in 100 bound protein complexes randomly selected from our independent test set. For each complex we first selected a point in the target protein assumed to be the center of the interface, and then attempted to recover the bound complex within the 100 binder proteins comprising the test set (Fig. 5d). A successful prediction means that a predicted complex with an interface Root Mean Square Deviation (iRMSD) of less than 5 Å relative to the known complex is included in a shortlist of the top 100 results. For comparison, we performed the same task using PatchDock^34^, which to our knowledge is one of the fastest docking programs available, and compared our runtime performance and number of recovered complexes. PatchDock was able to solve 40 complexes, with a mean iRMSD of 1.64 Å, and a running time of 2854 minutes. Using MaSIF-search with *K*=100 matches, we recovered 41 native complexes in 2 minutes (excluding database precomputation time), with mean interface RMSD of 2.36 Å. By increasing K to 3000 patches, we recovered 62 complexes in 17 minutes, with mean interface RMSD of 2.50 Å. Lastly, we compared the performance of our second stage alignment method coupled to GIF descriptors, this resulted in only 9 recovered complexes.

Even though we trained only on co-crystallized protein complexes, we also tested our method in a benchmark set of 40 proteins crystallized in the unbound (apo) state. Due to the fact that unbound docking is significantly more challenging, we increased the success criteria to 1 correct complex within the shortlist of the top 1000 results. PatchDock solved 11 complexes in 1045 minutes, while MaSIF-search solved 14 (K=1000) and 17 (K=3000) in 4 and 11 minutes, respectively; using GIF descriptors on apo proteins resulted in only 3 solved complexes.

In the previous test, we provided the site of the interface as input for docking; however, when the interface site is unknown, an attractive possibility is to combine MaSIF-site and MaSIF-search for the prediction of PPIs. To illustrate this strategy with a qualitative example, we selected the protein complex PD1:PD-L1^35^ (PDB id: 4ZQK). Assuming we did not have information about the PPI site in the PD1:PDL1 complex, we first used MaSIF-site for binding site prediction in the target protein PD-L1. This was followed by MaSIF-search to scan a database of ∼11,000 structures (52 million surface fingerprint descriptors) in order to find the best binders of PD-L1 (this protocol is shown in Supp. Fig. 5). The ground truth binder, PD-1 was included among the 11,000 structures. No PD1:PDL1 complexes were found in the training set. Our combined approach identified the mouse version of PD1 bound to human PD-L1 as the best binder (ranked #1, #3, #4), and the ground truth human PD-1 binder (ranked #8) in 26 minutes. Performing these tasks using traditional docking tools is prohibitively expensive, at tens of thousands of processing hours for even the fastest tools. In summary, MaSIF is able to decipher patterns that drive protein-protein interaction and encode them in a space that is amenable to perform fast searches. Thus, MaSIF provides an alternative to search vast protein databases for specific interaction fingerprints.

## Discussion

The molecular surface models the geometric and chemical features of a protein structure that contact other biomolecules. These features arrange in patterns, yet most of these patterns are difficult (if not impossible) to identify by human visual inspection, as they involve combinations of chemical and geometric features. MaSIF, thanks to the availability of large datasets and its deep learning framework, can help decipher them. This complexity is reflected in our results. As shown in our ablation tests, each input feature has a task-dependent effect on performance, and that highest performances are achieved by their combination. Thanks to our data-driven approach, these specific features may be learned for each task, which may be impossible with handcrafted approaches. This is exemplified in our MaSIF-search results, where we directly compared against handcrafted GIF descriptors, a previously presented technique for fast searches of protein surfaces. Our descriptors greatly outperform handcrafted GIF descriptors (Fig. 5d), consistent with the emergent trends in the field of computer vision and that could be critical to aid protein surface analysis.

One major advantage of a molecular surface description of protein structure is that it abstracts the underlying protein sequence. This abstraction may allow MaSIF to learn fingerprints that are independent of a protein’s evolutionary history. In each of our proof-of-concepts we performed a best-effort separation between training and testing datasets to test the capacity of MaSIF to learn sequence-independent interaction fingerprints, while still aiming to maintain large datasets for training and testing. For MaSIF-ligand we removed all homologous sequences (up to 30% identity). Even though 6 out of 7 metabolites have a common adenosine moiety, MaSIF was still able to decipher the pocket patterns that favor one metabolite over another, even in cases where the metabolites are highly similar. MaSIF-ligand exploits differences in both geometry and chemistry to determine the metabolite preference of proteins. To predict interaction sites (MaSIF-site) we performed both sequence and structural splits on the dataset, thus allowing us to predict evolutionary-independent fingerprints prone to protein-protein binding events. The predicted sites display non-obvious features that go beyond the typically recognizable surface patches displaying homogenous charge and hydrophobicity distributions. Finally, in MaSIF-search we encoded fingerprinted patterns, which enable ultrafast searches of potential binders that can be used for large-scale PPI prediction. Crucially, our general approach to learning interaction fingerprints may enable a more complete understanding of a protein’s function independently of its underlying sequence or other structural relationships. This may prove to be critical in fields of protein science that have been shifting away from the space of evolutionary-related proteins, such as *de novo* protein design.^36^ Moreover, even though we abstracted the sequence and a certain level of structural information (e.g protein fold) of proteins in an effort to understand the limits of learning from the surface, MaSIF has no intrinsic limitation to the supplementation of sequence, structure, experimental data, or other information additional to the learning process, and could serve as a complement to other tools that exploit these data sources.

Some important limitations arose during our experiments which can help to lead the way for the next-generation developments. MaSIF-search can be useful to search large databases for potential binding partners, yet its predictive power seems to depend significantly on the shape complementarity of the partners (Supp. Fig. 6). Thus, MaSIF-search’s ultrafast capabilities are likely to be useful mainly in cases where surfaces have high complementarity, and less so in highly solvated PPIs. In the future, biasing training sets towards more solvated interfaces, as well as more complex neural network architectures may help alleviate this issue. Furthermore, our training datasets consisted of proteins co-crystallized with their interaction partners, since these are more numerous and better annotated than *apo* proteins. When tested on *apo* structures, we see a deterioration of performance for *MaSIF-search*. In the field of computer vision, augmenting datasets with artificially generated data often leads to better neural network performance^37^. It is likely that augmenting MaSIF’s training datasets to incorporate simulated *apo* states using simulation tools (e.g., molecular dynamics) may improve the prediction on real *apo* proteins.

The proof-of-concept applications presented here meant to showcase MaSIF’s generality and the concept of learning from surface features, yet we believe they will still be useful to biologists. We foresee applications for MaSIF-ligand for proteins crystallized in the apo (unbound) state, which could be efficiently integrated with systems-level experiments^38^ for the characterization of large-scale ligand-protein interaction networks. MaSIF-site could provide “surface hot-spots,” which may be more easily targeted in the design of novel biologics for therapeutic purposes. MaSIF-search could be coupled to experimental or sequence-based method to identify potential binding partners for proteins, or it could be used to find potential engaging partners to use as starting points for protein design^29, 33^. Beyond these applications, we foresee our fingerprint-based approach to be particularly useful for the field of computational protein design. The computational design of new biomolecular interactions remains a fundamentally unsolved problem in protein design, despite notable advances. In the future, protein design programs such as Osprey^39^ and Rosetta^40^ may become fingerprint-aware, optimizing the sequence of *de novo*-designed proteins to display the interaction fingerprints on the molecular surface necessary to perform a functional task.

Collectively, we present a conceptual framework to decipher interaction fingerprints, leveraging the representation of protein structures as molecular surfaces, together with powerful data-driven learning techniques. The availability of our data and code will allow researchers to apply our framework to new problems. Our current applications show important technical advantages with great potential for further development and considerable impact on the fundamental study of protein structure and function, as well as for the design of novel proteins and protein-based therapies.

## Acknowledgements

The authors would like to thank Jaume Bonet, Sabrina Vollers, Paolo de los Rios and Sarel Fleishman for critical reading of the manuscript and helpful comments. This work was funded by generous grants from the European Research Council (Starting grant — <u>716058</u>), the Swiss National Science Foundation, Novartis Foundation for Medical-Biological Research and the Biltema Foundation to B.E.C. P. Gainza is sponsored by an EPFL-Fellows grant funded by an H2020 Marie Sklodowska-Curie action.

## Contributions

P.G., F.S., F.M., M.B., and B.E.C designed the overall method and approach. M.B. and B.E.C supervised the research. P.G. designed and implemented MaSIF-site and MaSIF-search. F.S. designed and implemented MaSIF-ligand. PG and FS developed MaSIF-search’s second-stage alignment algorithm. P.G., F.S., M.B. and B.E.C. analyzed the data. E.R. assisted in the design and development of these methods. P.G., F.S. and B.E.C wrote the manuscript. All authors read and commented the manuscript.

## Methods

### Computation of molecular surfaces

All proteins in the datasets were protonated using Reduce^41^, and triangulated using the MSMS program^42^ with a density of 3.0 and a water probe radius of 1.5 Å. Protein meshes were then downsampled and regularized to a resolution of 1.0 using pymesh^43^. Geometric and chemical features were computed directly on the protein mesh, with the exception of the distance-dependent curvature, which was computed on each patch according to the surface normals of the vertices in the patch.

### Decomposition of proteins into overlapping radial patches and computation of features

For each point in the discretized protein surface mesh, a radial patch of geodesic radius 9 Å or 12 Å (application-dependent) was extracted to perform an analysis of the surface features of the patch. The following features were included in each patch:

**Shape index** - describes the shape around each point on the surface, with respect to the local curvature [15]. Values range from −1 (highly concave) to +1 (highly convex). It is defined with respect to the *principal curvatures κ*_1_, *κ*_2_, *κ*_1_ ≥ *κ*_2_ as:

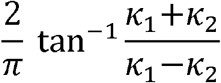

**Distance-dependent curvature** - for every vertex within an extracted patch, the distance-dependent curvature computes a value in the range [-0.7, 0.7] that describes the relationship between the distance to the center and the surface normals of each point and the center point. Details of this feature are described in ^15^. While the principal curvature component describes the shape around each vertex in the full protein, we found that it is also informative to compute the curvature within each patch, using the center of the patch as a reference.

**Poisson-Boltzmann continuum electrostatics** - PDB2PQR^44^ was used to prepare protein files for electrostatic calculations, and APBS^45^ (version 1.5) was used to compute Poisson-Boltzmann electrostatics for each protein. The corresponding charge at each vertex of the meshed surface was assigned using Multivalue, provided within the APBS^45^ suite. Charge values above +30 and below −30 were capped at those values and then values were normalized between −1 and 1.

**Free electrons and proton donors** - the location of free electrons and potential hydrogen bond donors in the molecular surface was computed using a hydrogen bond potential^22^ as a reference. Vertices in the molecular surface whose closest atom is a polar hydrogen, a nitrogen, or an oxygen were considered potential donors or acceptors in hydrogen bonds. Then, a value from a Gaussian distribution was assigned to each vertex depending on the orientation between the heavy atoms^22^. These values range from −1 (optimal position for a hydrogen bond acceptor) to +1 (optimal position for a hydrogen bond donor).

**Hydropathy** - Each vertex was assigned a hydropathy scalar value according to the Kyte & Doolittle^20^ scale of the amino acid identity of the atom closest to the vertex. These values, in original scale ranged between −4.5 (hydrophilic) to +4.5 (most hydrophobic) and were then normalized to be between −1 and 1.

### Computation of geodesic polar coordinates

Once surface patches are extracted from a protein, MaSIF uses a geodesic polar coordinate system to map the position of vertices in radial (i.e. geodesic distance from the center) and angular coordinates (i.e. angle with respect to a random directions) with respect to the center of the patch (Fig. 1c). These coordinates add information on the spatial relationship between features to the learning method.

**Radial coordinates** - describe the geodesic distance of a point to the center of the patch. Due to its speed, we used the Dijkstra algorithm implemented in Matlab to compute an approximation of the true geodesic distance.

**Angular coordinates** - to compute angular coordinates for each vertex in the grid, a classical multidimensional scaling algorithm^46^ implemented in Matlab was used to flatten patches into a plane based on the pairwise geodesic distances between all vertices. As molecular surface patches have no canonical orientation, a random direction in the computed plane was chosen as a reference, and the angle of each vertex to this reference in the plane was set as the angular coordinate.

### Geometric deep learning on a learned soft grid

Geometric deep learning allows us to apply successful image-based deep neural network architectures, such as convolutional neural networks^16^, to geometric data such as surfaces. Our approach relies upon establishing a mapping from the protein surface’s geodesic domain to a 2-D domain^17^. We first map a surface patch to a 2-D learned soft polar grid and then apply established deep learning layers, such as convolution^16^, on the mapped data. Our learned polar grid contains θ angular bins, and ρ polar bins, for a total of *J* - ρ *θ* bins. For each vertex in the discretized molecular surface *x*, with neighbors *N*(*x*), and each vertex *y* ∈ *N*(*x*), we define the coordinates *N*(*x*, *y*), the radial and angular coordinates of *y* with respect to *x*. The mapping of each grid cell *j* for feature vector *f* and the patch centered at *x*, *D_j_*(*x*) *f* is defined as:

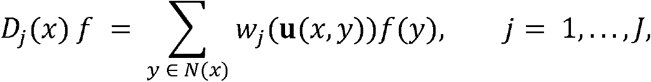

where *w_j_* is a weight function, and *f*(*y*) are the features at vertex *y*.

**Rotation invariance** - Rotation invariance is handled in the neural network by performing θ rotations of the input patch and performing a max-pool operation on the output^16^.

### MaSIF-ligand - ligand site prediction and classification

**Dataset** - All structures in the Protein Data Bank (PDB; 16 Oct 2018) including a protein chain but no DNA or RNA were considered if they included any of these seven chemical identifiers: ADP, COA, FAD, HEM, NAD, NAP or SAM. This resulted in 1853 ADP structures, 490 COA, 2020 FAD, 4448 HEM, 1269 NAD, 1212 NAP and 393 SAM. After building the biological assembly of these structures, the dataset was filtered based on sequence identity, to reduce redundancy and similarity between structures in the training and test sets. The filtering was performed as follows: the PDB provides pre-computed sequence clusters based on 30% sequence identity; each protein structure in the dataset was associated with one or more of these clusters based on its protein chains; two protein structures were defined as homologous if the associated clusters of both proteins coincided. The dataset was then reduced by randomly sampling structures from the dataset, one at a time, while continuously eliminating their homologs from the sampling pool. This process resulted in a total of 1459 structures, which were then randomly assigned to training (72%), validation (8%) and testing (20%) sets. The surfaces for these structures were generated as described above, and patches of 12 Å radius extracted. If a the center point of a patch was less than 3 Å from an atom for any of the seven ligands, the patch was labeled as a part of the binding pocket of the corresponding ligand.

**Neural network architecture, cost function and training optimization** - The training step and network architecture was as follows: 32 patches (limited for memory reasons) were randomly sampled from a single binding pocket. Each patch was used as input in a network and mapped to a learned soft grid with 16 angular bins and 5 radial bins. Each feature type (2 geometric and 3 chemical features) was run through a separate neural network channel, where the learned soft grid layer was followed by a convolutional layer with 80 filters, an angular max pooling layer with 16 rotations, a rectified linear, and a fully connected layer. A fully connected layer then combined the output from each channel, and output to an 80-dimensional fingerprint. The resulting 32 fingerprints were multiplied together to generate an 80×80 covariance matrix. The architecture for this network is shown in Supp. Fig. 7. The covariance matrix was flattened and fed first to a 64-unit, fully connected layer with rectified linear activation, and then to a 7-unit, fully connected layer with linear activation, followed by a softmax cross-entropy loss. The network was trained for 20000 iterations with the Adam optimizer with a learning rate of 1E^-4^. The validation error was evaluated every epoch and the best network was selected based upon this value. To obtain more stable predictions, each pocket was sampled 100 times and the resulting 100 predictions were averaged to obtain the final prediction.

**Visualization of patches important for NADP/NAD discrimination in Supp. Fig. 1**- From the NAD binding pocket of the dehydrogenase:NAD pocket (PDB id: 2O4C), 32 points were randomly sampled 10000 times and binding predictions made for each, giving 10000 predictions (7-dimensional vectors). For each prediction the prediction ratio NAD/NAP was computed. The predictions giving the top 90th percentile for this ratio were picked and the frequency of the patches behind these predictions were computed. The frequencies were normalized and overlaid on the protein surface. Same procedure was applied for the dehydrogenase:NADP complex (PDB id: 2YJZ) except that the NAP/NAD ratio was computed.

### MaSIF-site - protein interaction site prediction

**Datasets** - The PRISM database^47^ of PPIs, a compendium of non-redundant PPIs found in crystal structures, was used as the first source. Proteins with parsing problems or that failed to complete the feature computation were discarded. The PRISM database contains many complexes formed by the contacting protein chains found in asymmetric crystal units, which likely do not form in solution. To remove those complexes, we discarded PPIs that have no pairs of patches below a minimum threshold of radial shape complementarity (set at S=0.4; see below for a definition). In total, 8466 proteins engaged in PPIs were taken from the PRISM database. In addition, 3536 non-obligate (transient) interactions were taken from three databases: the PDBBind database^48^, the SAbDab antibody:antigen database^49^, and the ZDock benchmark database^50^. Finally, the resulting 12002 proteins were clustered according to sequence identity using the psi-cd-hit^51^, and one representative member was chosen from each cluster, resulting in 3362 proteins. A pairwise matrix of all TM scores for these proteins was then computed, and a hierarchical clustering procedure using scikit-learn^52^ was used to split the sets, resulting in a training set of 3004 proteins and a testing set of 358 proteins.

**Definition of interface points in a protein surface** - We defined the ground truth interface as the region of the surface that becomes inaccessible to solvent molecules upon complex formation. This was done by computing the surfaces of the complexes and the unbound partners. Surface regions in the individual partners that have no corresponding surface in the bound complex were then defined as the ground truth interface. Surface regions that become solvent inaccessible upon complex formation were defined as the ground truth interface.

**Neural network, cost function, and training optimization** - A neural network with three convolutional layers was used for this application. A diagram of the architecture is shown in Supp. Fig. 8. The network received as input a full protein decomposed into overlapping surface patches with a radius of 9.0 Å. The patches are mapped onto learned grids with 3 radial bins and 4 angular bins. The output of the network is an *interface score* between 0 and 1 for each point in the surface. During training, the batch size consisted of a single protein, and the network was optimized using using an Adam optimizer^53^ on a sigmoid cross-entropy loss function. As the number of non-interface points is usually much larger than the number of interface points, a random subset of non-interface points was selected to train on an equal number of positive and negative samples.. The neural network was trained during 40 hours.

**Comparison to SPPIDER** - The performance of a set of 59 known transient interactions for the test set was compared with that of the interface predictor SPPIDER^27^. Each protein was uploaded to the Sppider web site (http://sppider.cchmc.org/) and a regression-based prediction was computed on each residue. Following SPPIDER’s definition of ground truth interface residues^27^ as closesly as possible, the ground-truth interface residues were defined as those whose solvent excluded surface changes at least 5 Å^2^ upon binding and at least 4% change in solvent accessibility. We note that we used the solvent-excluded surface for these calculations and not the solvent accessible surface. In order to perform a comparison with MaSIF-site, our predictions were converted to a per-residue scoring by assigning the maximum score of all the residue’s points in the surface. A comparison point-by-point is shown in Supp. Fig. 3a.

### MaSIF-search - prediction of PPIs based on surface fingerprints

**Datasets** - a dataset of protein pairs that were co-crystallized and shown to engage in PPIs were taken from the PRISM database (see above). In addition, 3536 non-obligate (transient) PPIs were taken as was done for the interface site prediction, forming a set of 6001 PPIs. The PPI structural interface was extracted from the native complexes and a pairwise TM-align^54^ score matrix with all interfaces was computed. Then, a hierarchical clustering of the structures was performed according to the TM-align score using scikit-learn’s hierarchical clustering (AgglomerativeClustering)^52^. In total, the dataset was split into 4944 training PPI pairs and 957 testing PPIs. This list is complemented by 40 apo complexes, corresponding to those proteins in the testing PPIs such that both partners’ *apo* crystal structure was available in the Zdock benchmark^50^. The list of PDBs in the training and testing sets are provided in our github repository.

**Selection of interacting and non-interacting patches** - For each PPI, all pairs of surface patch centers belonging to distinct proteins and within 1.0 Å of each other were considered further. A *radial shape complementarity* score was computed for the pair as follows: (i) the shape complementarity of each point in the patch to the neighboring patch was computed; (ii) points within 12 Å of the center were divided into 10 concentric radial bins, in increments of 1.2 Å; the shape complementarity of the bin was computed as the 25^th^ percentile of the points in the bin; (iii) the radial shape complementarity S of the patch was computed as the median across all bins. The neural network for Fig. 5 was computed with interacting patches with a value of S > 0.5, while different ranges of S (−1 < S < 0.1 for very low complementarity, 0.1 < S < 0.3 for low complementarity, and 0.3 < S < 1.0 for high complementarity) were also tested and are shown in Supp. Fig. 6. Non-interacting pairs were selected by pairing a truly interacting patch with a randomly chosen one from any other protein in the set.

**Neural network architecture, cost function and training optimization** - The Masif-search neural network receives the features of one patch (which may be inverted for the binding partner) as input and then outputs a vectorized descriptor. The architecture for this network is shown in Supp. Fig. 9. During training and testing, a *binder*, a *target* and a random patch are always input into the network, such that the binder and target are known interacting pairs, and the target and random are assumed to be non-interacting. The features for the target are inverted (multiplied by −1), with the exception of the hydropathy index. A total of 85652 true interacting pairs and 85652 non-interacting pairs were chosen for training/validation, while 12678 true interacting and 12678 non-interacting pairs were chosen for testing. The network was trained to minimize the Euclidean distance between the fingerprint descriptors of binder and target, and maximize the distance between the descriptors of target and random. Each patch was input to a network and mapped to a learned soft grid with 16 angular bins and 5 radial bins. Each feature type (2 geometric and 3 chemical features) was ran through a separate neural network channel, where the learned soft grid layer was followed by a convolutional layer with 80 filters, an angular max pooling layer with 16 rotations^18^, and a rectified linear unit. A fully connected layer then combined the output from each channel, and output an 80- dimensional fingerprint. During training, each iteration batch consists of a set of 8 binder/target interacting pairs and 8 target/random interacting pairs. The optimization process during training, using an Adam optimizer^53^, consists of minimizing the d-prime cost function^55^:

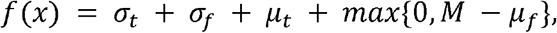

where *µ*and*µ* are the median distance for true and non-interacting pairs, respectively, while *σ*and *σ*are the standard deviation for true and false interacting pairs. The neural network was trained with batches consisting of 8 binder, 8 target, and 8 random patches. The network was trained for 40 hours.

**Structural alignment and rescoring** - A second-stage alignment and scoring method generates the complexes based on the identified fingerprints. The top K patches with the shortest fingerprint descriptor distance to the target patch are selected as a short list of potential binding partners. Each binder patch is then aligned using the RANSAC algorithm implemented in Open3D^56^ (Supp. Fig. 4). Briefly, RANSAC selects three random points from the binder patch and uses the computed descriptors to find the closest points in the target patch by descriptor distance. Using these three newly found correspondences, RANSAC attempts to align the source patch to the target patch. RANSAC iterates for either 2000 transformations (for the co-crystal benchmark) or 4000 transformations (for the apo state complex). The transformation is scored according to the number of points within 1.0 Å between binder and target. After RANSAC completes, the transformation is rescored according to the function 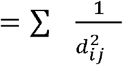 where is the descriptor distance between binder point and target points, such that and are within 1.0 Å. Once all K selected patches have been aligned, we rerank the list using score (Supp. Fig. 3). To optimize speed, the extracted patches were reduced to 9 Å.

**PPI search docking benchmark** - N=100 co-crystal structure complexes were selected from the testing set. One of the two proteins was selected as the target protein; for each target protein, the patch with the highest radial shape complementarity to the binder protein patch in the co-crystal structure was selected as the target site (Fig. 5d). The benchmark consisted of recovering the conformation of the binder within a short list of the 100 top-ranked results. A second benchmark was performed with N=40 complexes in the apo state, aligned to the bound complex. The benchmark for *apo* structures was performed in the same way as for the co-crystal structures, but the success criteria were relaxed to recovering the conformation of the binder within the top 1000 results. For all methods benchmarked, the binder was randomly rotated before performing any alignments.

**Comparison to GIF descriptors** - Geometric invariant fingerprint (GIF) descriptors were implemented to our best efforts according to the description by Yin *et al*^15^. For testing of the descriptors, the features of the target were inverted before computing the GIF descriptor. In the PPI search benchmark, GIF was coupled to our second stage alignment and scoring method.

**Comparison to PatchDock** - PatchDock^34^ was used with default settings, assigning the residue closest to the target site as an active site residue. After all alignments, PatchDock’s transformations for all N proteins were merged and scored according to PatchDock’s default Geometric Score.

**PDL1 benchmark**. A modified version of our second-stage alignment and scoring method was run on the PD-L1:PD1 complex found on the Protein Databank structure with PDB id: 4ZQK^35^. The exercise consisted on recovering the bound PD-L1:PD1 complex, among all possible complexes between PD-L1 and 10954 other proteins. First, the binding site scores on the surface of the PD-L1 chain (chain A in PDB id 4ZQK) was predicted using MaSIF-site. Then, the center of the interface was predicted by finding the patch with the highest mean interface score. Once the center of the interface was identified, the descriptor of this center point was matched to all patches in the 10954 proteins, for a total of 52 million fingerprints. Matches were ignored if the descriptor distance was greater than 1.7 or if the interface score was less than 0.9. The matches that passed this filter were explicitly aligned and reranked using our second stage alignment and rescoring protocol. The top ten matches were then visually identified, showing the mouse PD1 (PDB id: 3BIK) as the top scoring transformation, followed by the real, wildtype match ranked #8.

### Code availability

Code availability - All code was implemented in Python and Matlab. Neural networks were implemented using TensorFlow. Both the code and scripts to reproduce the experiments of this paper are available at: https://github.com/lpdi-epfl/masif. The bound PDBs in the training/testing set and the computed surfaces with chemical features are available at Zenodo with DOI: 10.5281/zenodo.2625420. The unbound PDBs in the test set are provided in the github repository. We also provide a PyMOL plugin for the visualization of these structures.

**Pre-computation and neural network running times** - The time it takes to precompute the data and run the neural network on a 100 amino acid protein structure is approximately 2 minutes. Data precomputations were performed on CPUs, while neural network calculations were performed in GPUs.

---

**Supp. Fig. 1.**
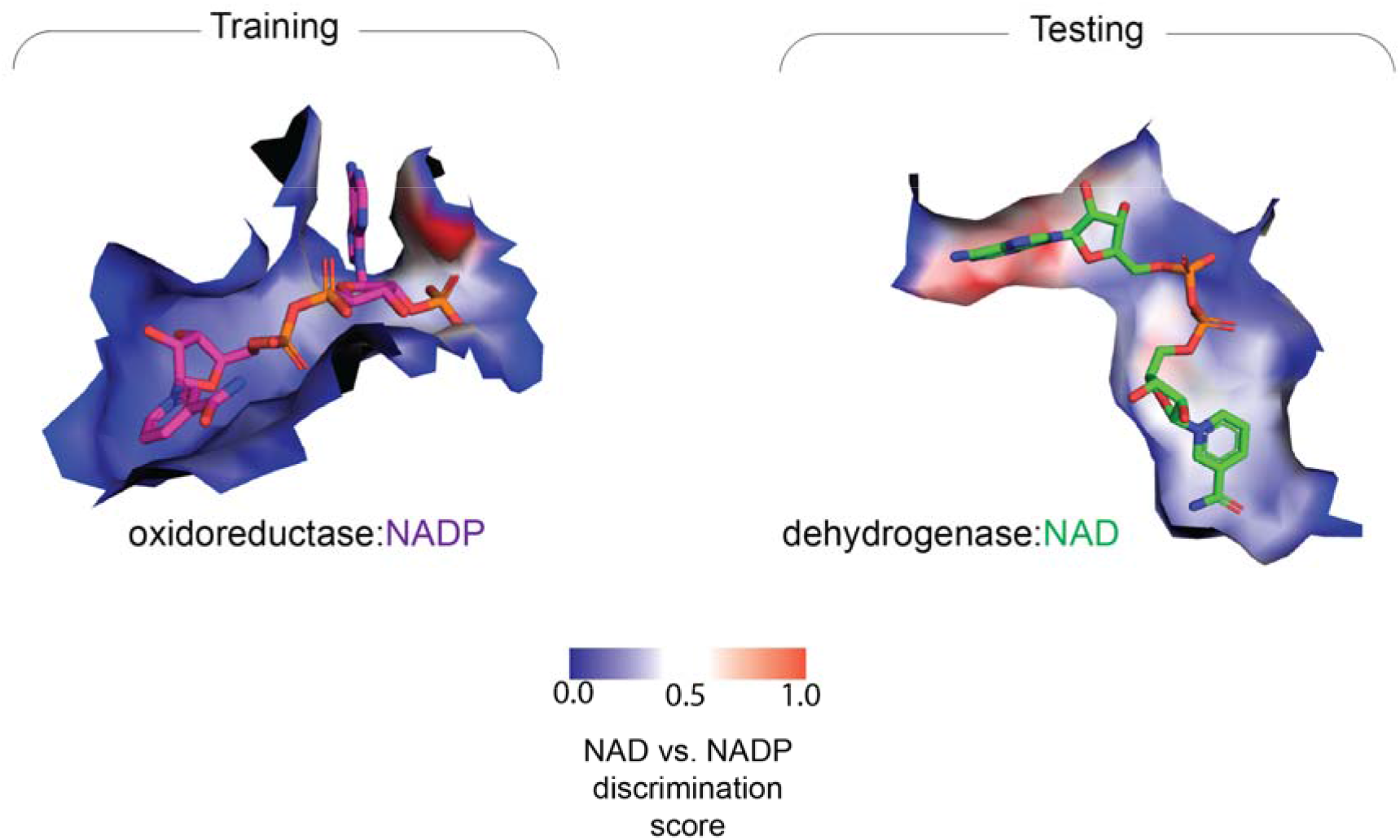
Analysis of MaSIF-ligand’s discrimination between NADP and NAD on two specific examples: a bacterial oxidoreductase and a human dehydrogenase. The bacterial dehydrogenase in the test set binds to NAD (PDB id: 2O4C), while its closest structural homologue in the training set corresponds to a mammalian oxidoreductase (PDB id: 2YJZ)^25^, which binds to NADP^24^. Here we scored the pocket surface by a discrimination score, which scores each point in the protein surface by its weight in the neural network’s distinction between NADP and NAD. Surface regions with high importance are shown in red, while those of low importance are shown in blue.

**Supp. Fig. 2.**
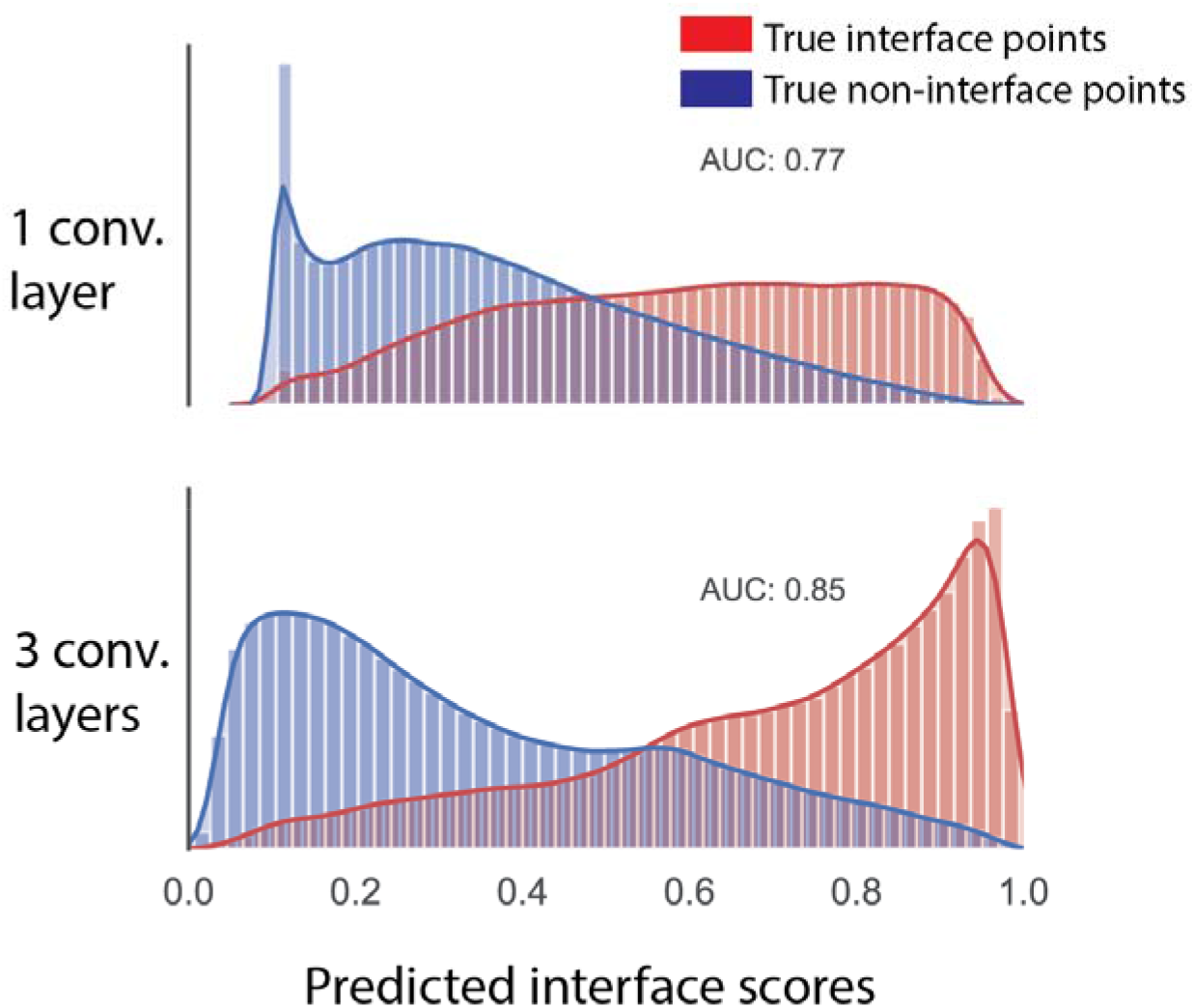
MaSIF-site interface site prediction predicted scores distribution for true positives (red) vs. true negatives (blue) with one convolutional layer (top) and multiple convolutional layers (bottom).

**Supp. Fig. 3.**
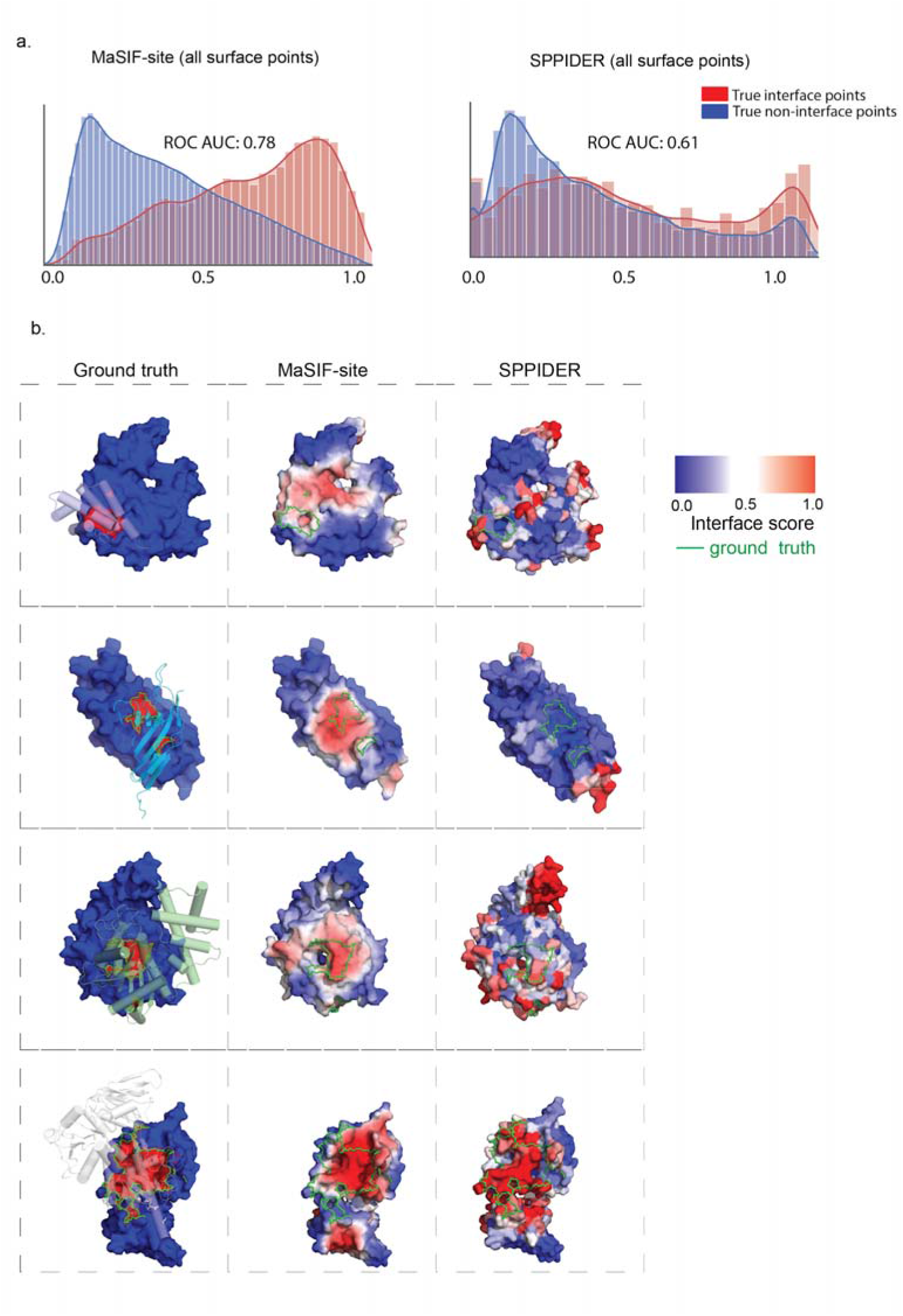
Comparison between MaSIF site and SPPIDER on transient interactions. **a**. Evaluation of MaSIF-site vs. SPPIDER on 59 transient interactions on a point-by-point basis. **Left:** Distribution of predicted interface points for true and false interface points for MaSIF-site. **Right:** Distribution of predicted interface points for true and false interface points for SPPIDER^27^. **b**. Randomly-selected examples from the testing set comparing our prediction with SPPIDER.

**Supp. Fig. 4.**
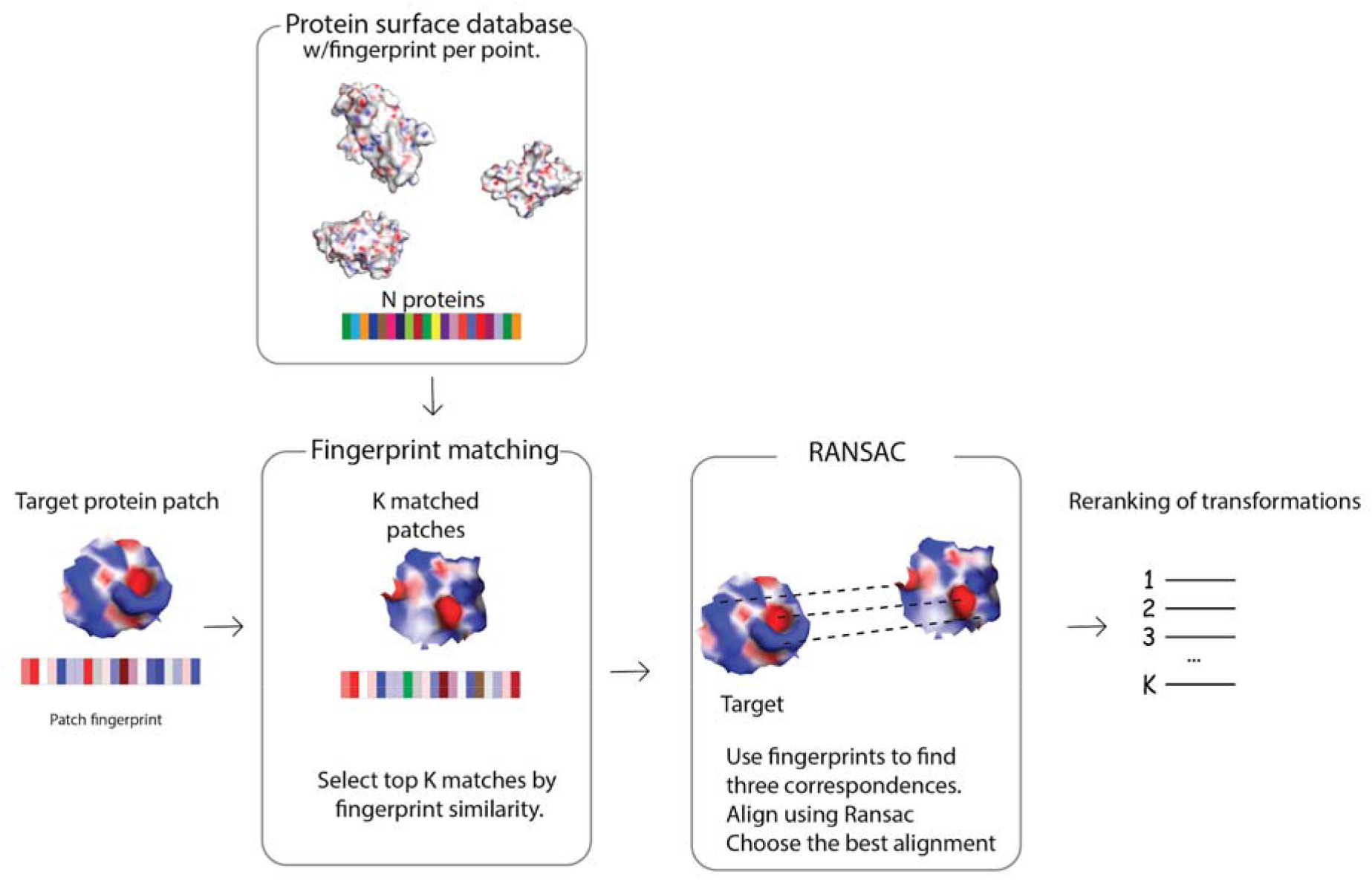
The MaSIF-search second-stage protocol. A fingerprint is computed on a selected target site (left). A database of hundreds of thousands of proteins with a fingerprint per point is searched for the K-most similar descriptors. Once these descriptors are matched, a set of correspondences between the matched patches is found using the RANSAC algorithm, and a transformation is obtained. Finally the transformations are reranked based on descriptor distances of all contact points (see Methods).

**Supp. Fig. 5.**
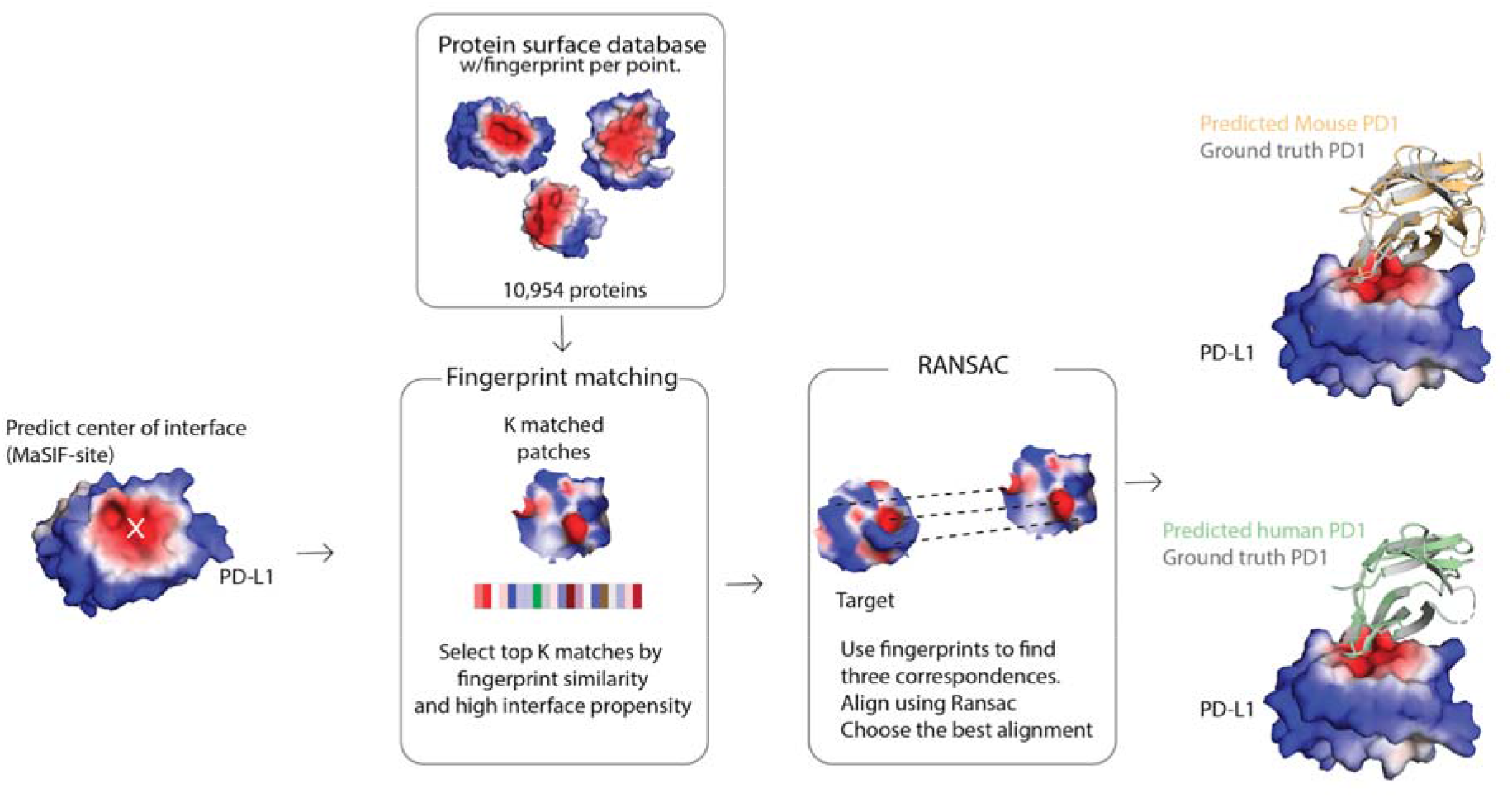
Hybrid MaSIF-search/MaSIF-site protocol to identify true binders in this example against PD-L1. The target site is first predicted using MaSIF-site. Then a database of nearly 11,000 proteins is scanned, and all patches with a MaSIF-site score > 0.9, and with a descriptor distance less than 1.7 are selected for alignments. Top candidates are matched using RANSAC, and reranked using the descriptor distance of all aligned points (described in Methods). The top predicted complex was the PD-L1:Mouse PD1 (PDB id: 3BIK), ranked #1 with an RMSD of 0.6Å (shown here in pale orange). The PD-L1:Human PD1 (PDB id: 4ZQK), was ranked #8 with an RMSD of 0.3 Å. Both are shown overlaid over the initial complex (PDB id: 4ZQK). The entire runtime protocol took approximately 26 minutes (excluding descriptor precomputation time).

**Supp. Fig. 6.**
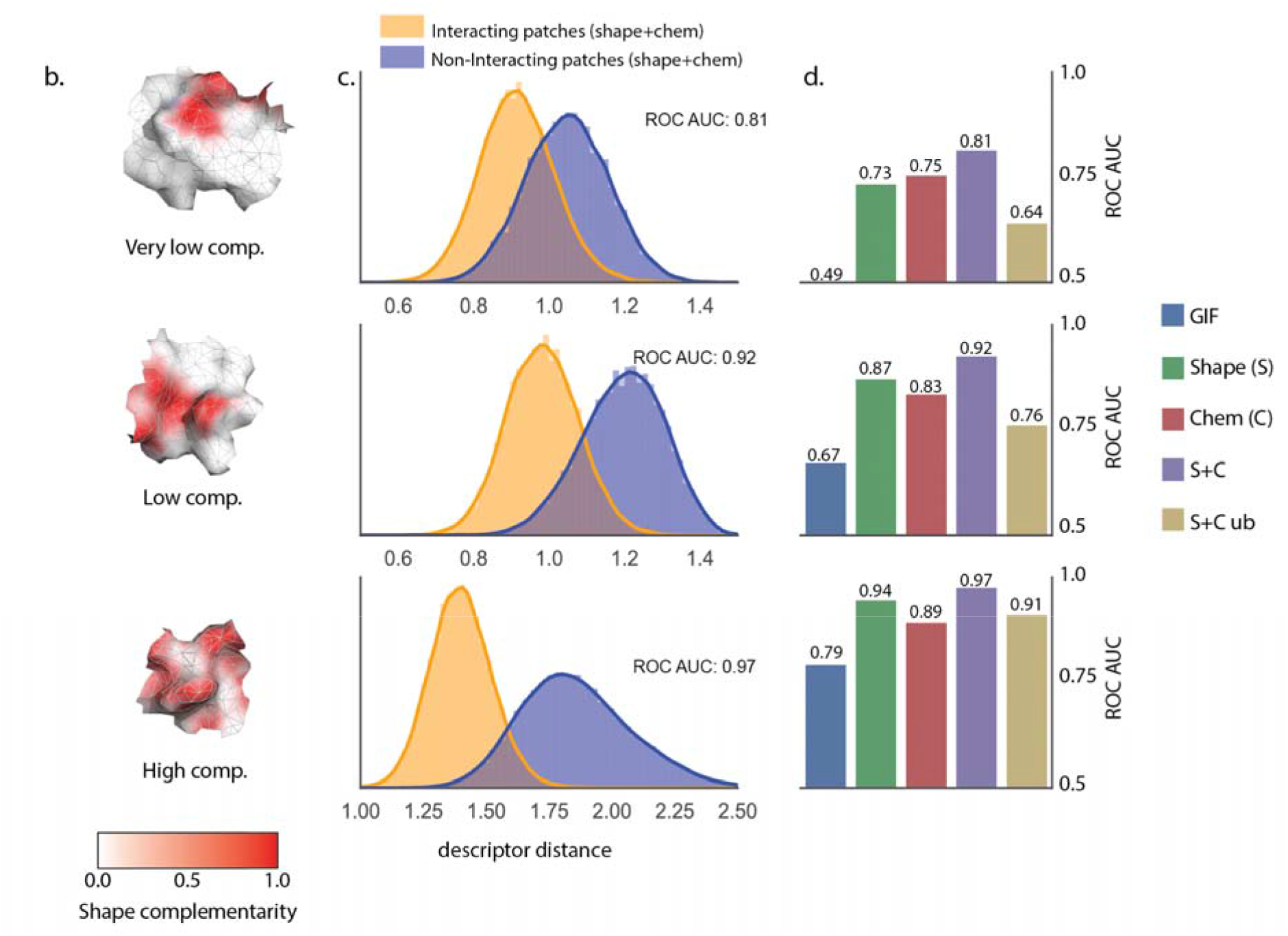
Performance of MaSIF-search fingerprints under different shape complementarity filters for the interacting patches. **a**. We set up three classes of interacting patches, filtered by shape complementarity, illustrated here with three examples, where the surface is colored according to shape complementarity from white (0.0) to red (1.0). **b**. Descriptor distance distribution plot for interacting and non-interacting patches depending on the shape complementarity class. **c**. ROC AUC values for the GIF descriptors^15^, descriptors trained only on geometry, chemistry, or both, and patches found in unbound protein within each complementarity class (S+C ub).

**Supp. Fig. 7.**
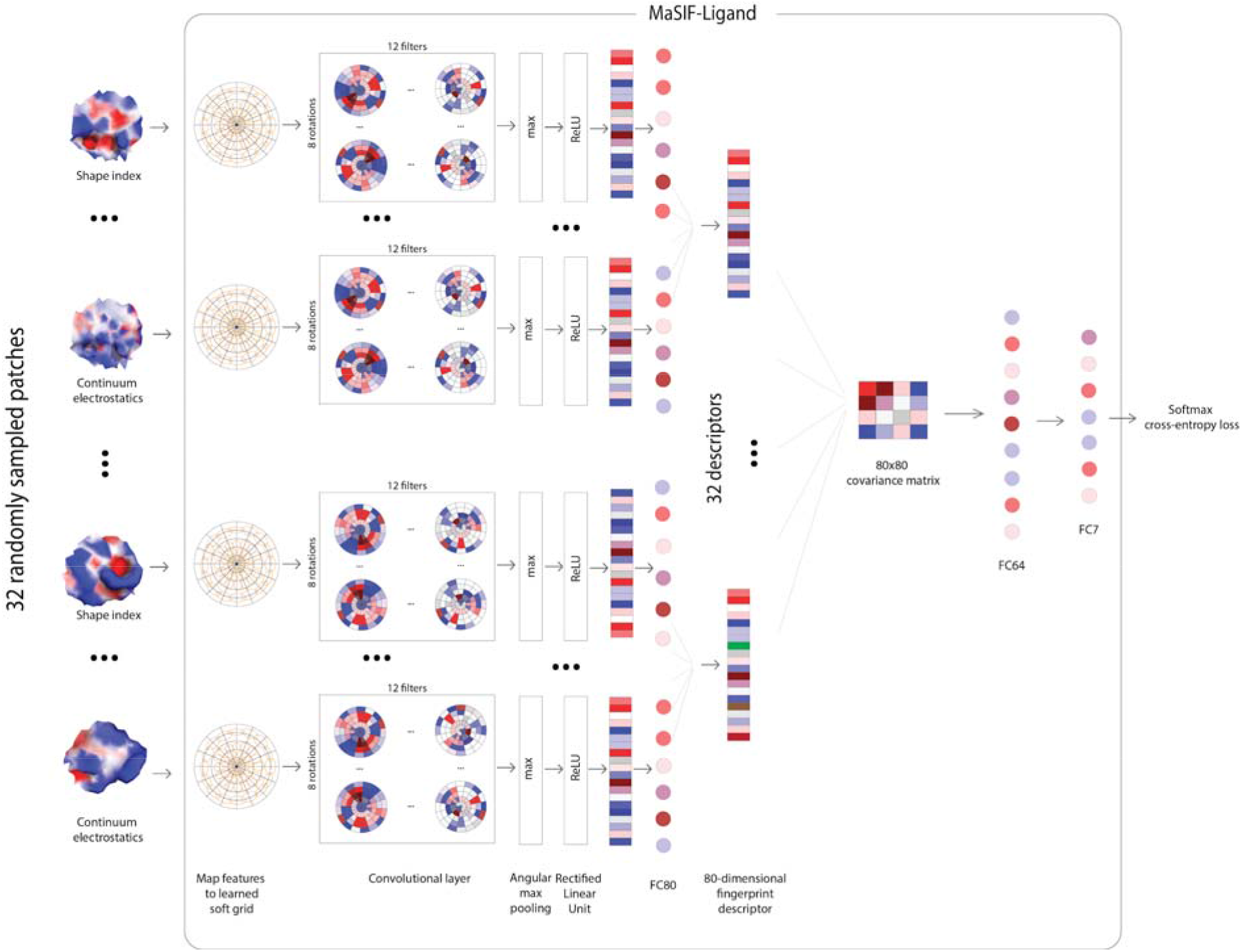
Network architecture for MaSIF-ligand. 32 randomly sampled pocket patches are fed through convolutional layers followed by a fully connected layer (FC80). Descriptors are combined in a 80×80 covariance matrix followed by two fully connected layers (FC64 and FC7) and then softmax cross-entropy loss.

**Supp. Fig. 8.**
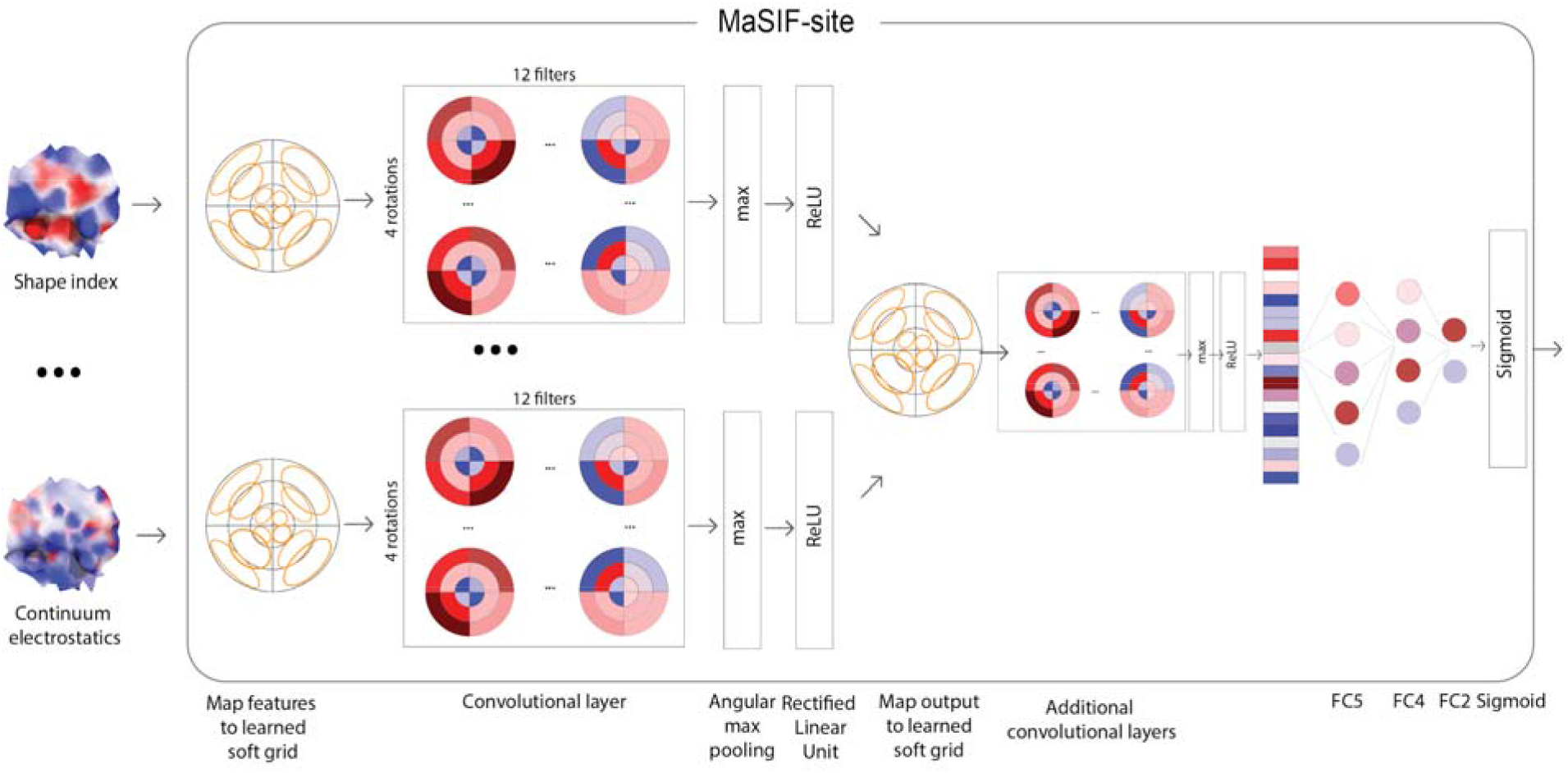
Network architecture for MaSIF-site. Patches are fed through convolutional layers followed by a series of fully connected layers (FC5, FC4, FC2), and finally a sigmoid cross-entropy loss.

**Supp. Fig. 9.**
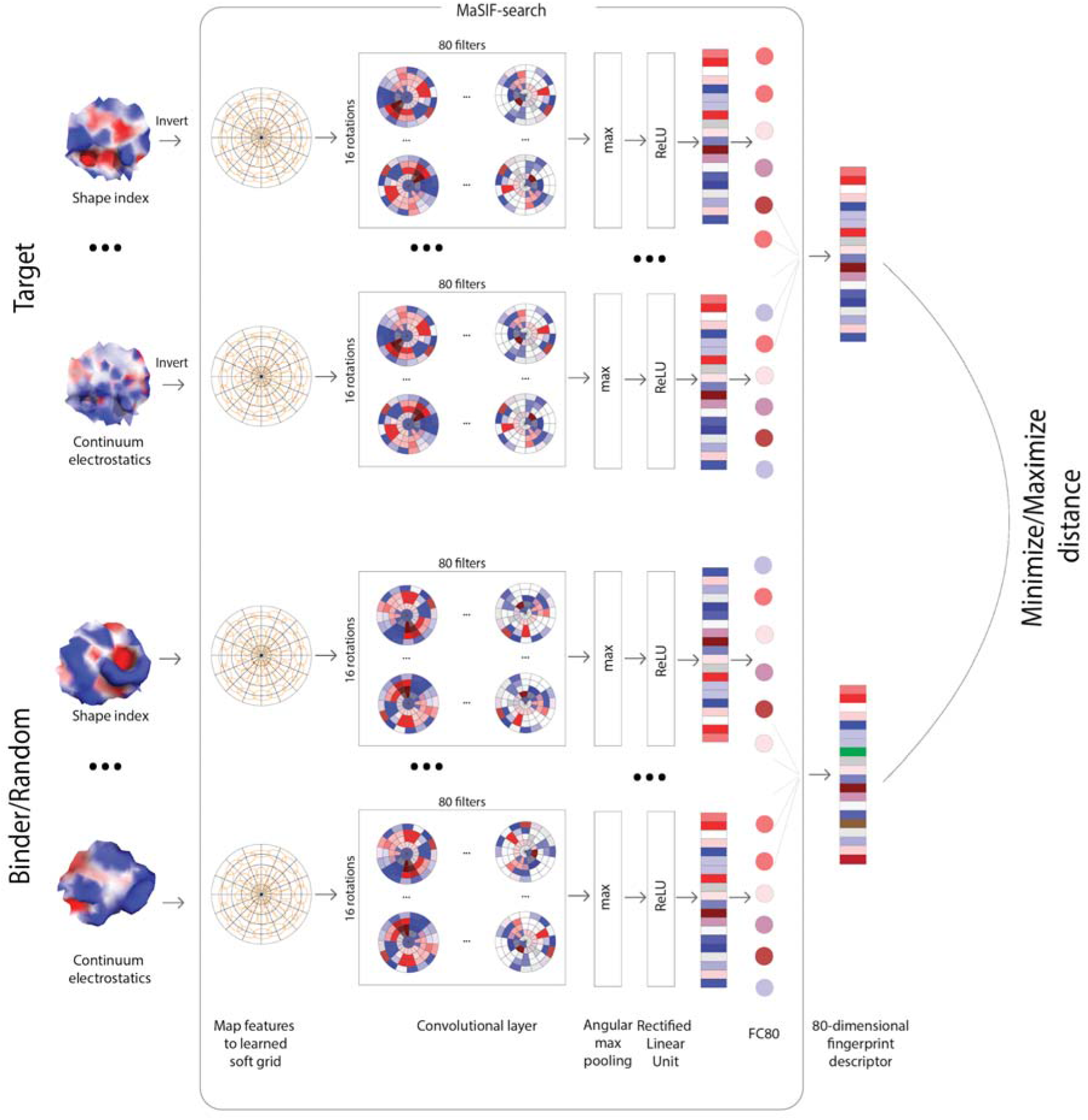
Network architecture for MaSIF-search. Patches from the target and the corresponding binder or a random patch are fed through convolutional layers, followed by a fully connected layer (FC80). The L2-distance between the resulting descriptors is computed and the neural network is optimized to minimize this distance with respect to binder and maximize it with respect to the random patch.

